# Expression of recombinant cardosin B in tobacco BY2 cells: an alternative system for the production of active milk clotting enzymes

**DOI:** 10.1101/2021.02.23.432499

**Authors:** André Folgado, Rita Abranches

## Abstract

*Cynara cardunculus* L. or cardoon is a plant that is used as a source of milk clotting enzymes during traditional cheese manufacturing. This clotting activity is due to aspartic proteases (APs) found in the cardoon flower, named cyprosins and cardosins. APs from cardoon flowers display a great degree of heterogeneity, resulting in variable milk clotting activities and directly influencing the final product. Producing these APs using alternative platforms such as bacteria or yeast has proven challenging, which is hampering their implementation on an industrial scale. We have developed tobacco BY2 cell lines as an alternative plant-based platform for the production of cardosin B. These cultures successfully produced active cardosin B and a purification pipeline was developed to obtain isolated cardosin B. The enzyme displayed proteolytic activity towards milk caseins and milk clotting activity under standard cheese manufacturing conditions. We also identified an unprocessed form of cardosin B and further investigated its activation process. The use of protease-specific inhibitors suggested a possible role for a cysteine protease in cardosin B processing. Mass spectrometry analysis identified three cysteine proteases containing a granulin-domain as candidates for cardosin B processing. These findings suggest an interaction between these two groups of proteases and contribute to an understanding of the mechanisms behind the regulation and processing of plant APs. This work also paves the way for the use of tobacco BY2 cells as an alternative production system for active cardosins and represents an important advancement towards the industrial production of cardoon APs.

## Introduction

Traditional cheeses in countries worldwide are produced using plant extracts as rennet to clot sheep and goat milk during cheese manufacturing ^1^. Although these products usually constitute a niche market, they represent an important source of income for local communities and also contribute to the conservation and valorization of those specific animal breeds used in manufacturing these products ^2^. High production costs are partly due to the provenance of the rennet from the flowers of certain plant species. The flowers of *Cynara cardunculus* L. or cardoon are commercially sold, mainly in the Iberian Peninsula, for their use as a milk clotting agent in the first step of traditional cheese manufacturing. This clotting activity is due to the presence of aspartic proteases (APs) in the cardoon flowers ^3,4^. However, a high level of heterogeneity among the flowers can sometimes lead to unpredictability in the cheese manufacturing process, ultimately leading to unsuccessful clotting ^5^.

To date, eleven different cardoon APs, named cardosins and cyprosins, have been identified at either the protein or RNA levels in the cardoon flower, with a high degree of similarity among them ^4–7^. All of these APs are typical of the A1 plant family. Like other APs, cardoon APs are synthesized as an inactive enzyme known as zymogen composed of a signal peptide, a prosegment involved both in the activation and folding of the protein, plus a 100-amino acid segment between the N-terminal and the C-terminal domains of the protein, which is called a plant specific insert (PSI) and is only found in plant APs ^8^. Its function is not yet fully understood, but evidence exists of its involvement in intracellular sorting and plant defense mechanisms ^9,10^. Before becoming active, plant APs undergo sequential proteolytic steps to remove certain sequence segments (namely the prosegment and the PSI) ^11^ leaving two segments, a light- and a heavy-chain linked by hydrophobic interactions and hydrogen bonds (Fig. S1) ^12^.

Attempts to produce some of these proteases in heterologous systems have not yet proven viable. The expression of cyprosin B in *Pichia pastoris* ^14^ and *Saccharomyces cerevisiae* ^15^ showed differing results in the two systems. In the former, processed cyprosin B accumulated in the culture medium, without completely removing the PSI, resulting in the heavy and light chains of the protein held together by disulfide bonds. This differs from the native cyprosin B and led to distinct structures and catalytic efficiencies ^13^. More recently, a *Galium verum* L. AP was expressed in *Pichia pastoris* ^14^ and analysis of the growth curve revealed protein secretion in its unprocessed form which was later processed due to pH fluctuations in the culture medium. The medium at some point became acidic, and the secreted unprocessed form underwent processing ^14^. Expression in *Saccharomyces cerevisiae* showed that recombinant cyprosin B was secreted into the culture medium in both its active and inactive forms, although with limited secretion and significant quantities remaining within the cells ^15^. In another report, synthetic cardosin B was produced and secreted in *Kluyveromyces lactis* ^16^. In this work, the PSI was removed and the two subunits were linked with glycines, which enabled the production and secretion of an inactive form of the enzyme which then required activation ^16^.

The viability of plant systems as a cardoon AP source has also been assessed. Cyprosin B was produced by genetic transformation of cardoon callus in order to overexpress the protein and characterize production in a bioreactor. Active cyprosin B was obtained, but the recombinant protein was accumulated inside the cells and not secreted to the medium. The protein was purified but yielded small quantities, which limited industrial scale use ^18^. More recently, cardoon hairy roots (HR) have been produced and characterized for their proteolytic content and were found to contain several APs among other protease classes ^17^. Milk clotting did take place; however, low quantities of cardosins and slow HR growth still limited the usefulness of this system as an alternative source of milk-clotting APs. These findings reveal that enzyme activation is highly dependent on the plant system, a fact which must be taken into account when selecting a cost-effective production platform.

This work investigated tobacco BY2 cell cultures as a plant-based platform for the production of cardosins. To our knowledge, this is the first time that this widely used system has been successfully applied for producing proteases for the food industry. We have demonstrated that BY2 cells can produce active cardosin B. Although recombinant cardosin accumulated within the cells, we developed an effective purification strategy. The purified cardosin B was then characterized regarding its activity on casein and milk-clotting. We also identified an unprocessed form of cardosin B and further investigated its activation process. We show that this system is suitable for producing active cardosins, thus paving the way for its industrial scale application.

## Results

### Recombinant cardosin B was retained intracellularly in BY2 cells

The literature reveals significant variation in cardosin expression patterns in cardoon flowers; cardosin A is found in the vacuoles of stigma cells, while cardosin B appears in the extracellular matrix of the style transmitting tissue. Due to these distinct locations and tissue natures, cardosin B is considered a secreted protein while cardosin A is defined as a vacuolar one ^18^. Our first goal was to investigate whether cardosin B follows the same route in BY2 cells as it does in the flower. To this end, the spent medium and cellular extract were analyzed by western blotting using a heavy-chain cardosin B specific antibody. As expected, no signal appeared in the spent medium and cellular extract of the WT line. However, in the cell lines expressing cardosin B, the protein was only detected in the cellular extract and not in the spent medium (Fig. 1a). This indicated that recombinant cardosin was not secreted, despite the fact that the vector contained a signal peptide.

**Figure 1.**
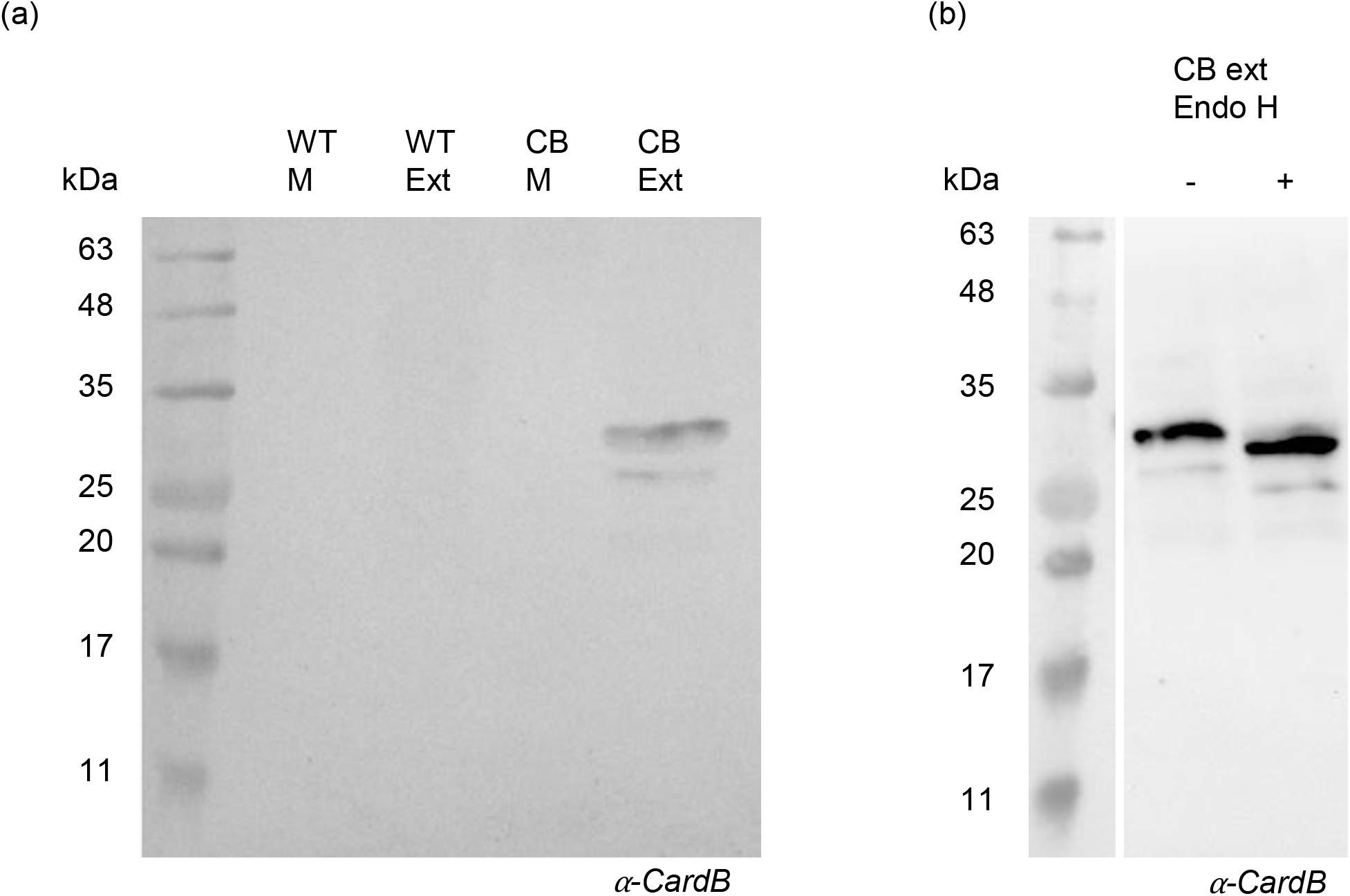
Detection of cardosin B heavy-chain by western blot. (**a**) Western blot of spent medium and cellular extracts of WT and CardB lines. (**b**) Endoglucosidase H assay using cellular extract of cardB lines.

In the cell extracts, two bands were detected, one sized 34 kDa, consistent with processed cardosin B ^4^ and a lower sized band that could be a different glycoform. To address this question, we performed an endoglicosidase H assay. The Endo H treatment reduced the size of both bands in the treated sample, suggesting some degree of protein deglycosylation (Fig. 1b). Since EndoH cannot cleave complex glycans, this finding also indicated that cardosin B produced in BY2 cells lacks complex glycosylation in agreement with results for cardoon flowers and other heterologous systems ^12^.

### Cardosin B was purified from BY2 cell extracts

Since cardosin B was found intracellularly, cellular protein extracts were used for purification. This process normally takes place at pH 3 ^4^, as acidity excludes most protein contaminants, producing an enriched cardosin extract while also preventing phenolic oxidation ^19^. To investigate the applicability of this strategy, we tested various pH values to determine whether the pH of the extraction buffers would influence cardosin B extraction from BY2 cells. Total protein content per gram of cells was found to decrease as pH became more acidic (Fig. S2a). The same samples were then used for western blotting to assess whether the reduction of total protein correlated with a decreasing amount of cardosin B in each extract. We found that, as the pH became more acidic, the relative amounts of cardosin B decreased. In fact, cellular extract at pH 3 contained about 10% of the cardosin B present in an extract performed at pH 8 (Fig. S2b). Despite the lower amount of cardosin B, a concentrated sample of BY2 extract at pH 3 contained fewer contaminant proteins and yielded a cardosin B enriched extract (Fig. S2c). Low pH extracts limited the number of proteins, including cardosin B, thus this procedure represents a tradeoff between purity and quantity. Therefore, the selected strategy to purify the recombinant protein was an acidic extraction followed by extract concentration and buffer exchange before the chromatographic procedure.

The recombinant cardosin B was purified using ion exchange chromatography, after unsuccessful attempts to use the 6xHis tag (Fig. S3). The concentrated sample passed through a Q-sepharose anionic exchange column and eluted in a NaCl gradient. A gradual increase of 5% of NaCl was able to separate most of the proteins, which allowed us to collect purified cardosin B in the 50% NaCl elution fraction (Fig. 2a). The pooled fraction appeared to contain only cardosin B, since no other protein appeared in gel staining (Fig. 2b). The SDS-PAGE gel showed the two previously detected bands, plus another one above 10 kDa. This band is likely to correspond to the light chain of the enzyme, although with a lower molecular weight than that previously described for the cardoon flower ^4^. Cardosin B has been described as comprising a heavy and a light-chain of 34 kDa and 14 kDa respectively, held together by hydrophobic interactions and hydrogen bonds ^12^. Although shorter than that observed by Verissimo and colleagues, the size of the putative light-chain corresponded to the *in silico* estimate of the cardosin B light-chain tagged with 6xHis (11.2 kDa). To determine if the differing molecular weights of the two higher bands and light-chain were due to protein truncation, the three bands were extracted from the gel and analyzed independently using Triple TOF-MS. Although the fragments did cover all the expected sequence, it was found that none of the bands had undergone truncation at either the N- or C-termini that might yield a smaller sequence.

**Figure 2.**
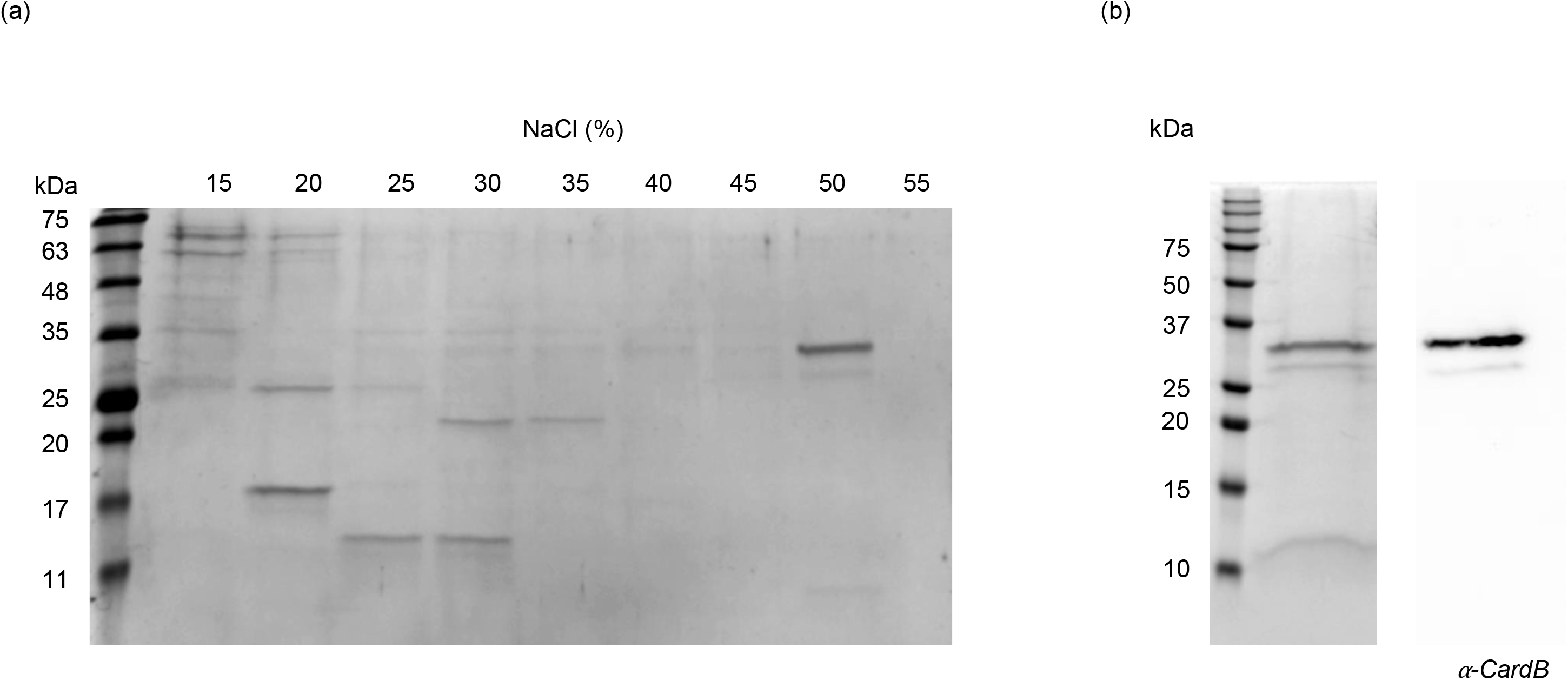
Purification of cardosin B by ion exchange chromatography. (**a**) SDS-PAGE analysis of NaCl gradient eluted fractions of cardosin B purification by Q-sepharose column (**b**) SDS-PAGE and western blot analysis of purified cardosin B. The molecular weight ladder used in figure A was NZYColour Marker II (NZYTech) while Precision Plus Protein WesternC Standards (Bio-Rad) was used in figure B.

### Recombinant cardosin B showed maximum activity at pH 5.5 and 50 °C and displayed milk clotting activity

To further study the activity of the purified cardosin B, proteolytic activity was assessed using azocasein as substrate, and optimal pH and temperature conditions were determined. Azocasein solutions were used with a pH range of 4.6 to 8 and incubated at 37 °C. The purified enzyme showed maximum activity at pH 5.5 and none at pH 8 (Fig. 3a). Optimal temperature was determined using an azocasein solution at pH 5.5 and temperatures ranging from 30 °C to 75 °C. The results showed enzymatic activity at a broad range of temperatures with optimal activity at 50 °C, with a marked decrease at higher temperatures (Fig. 3b).

**Figure 3.**
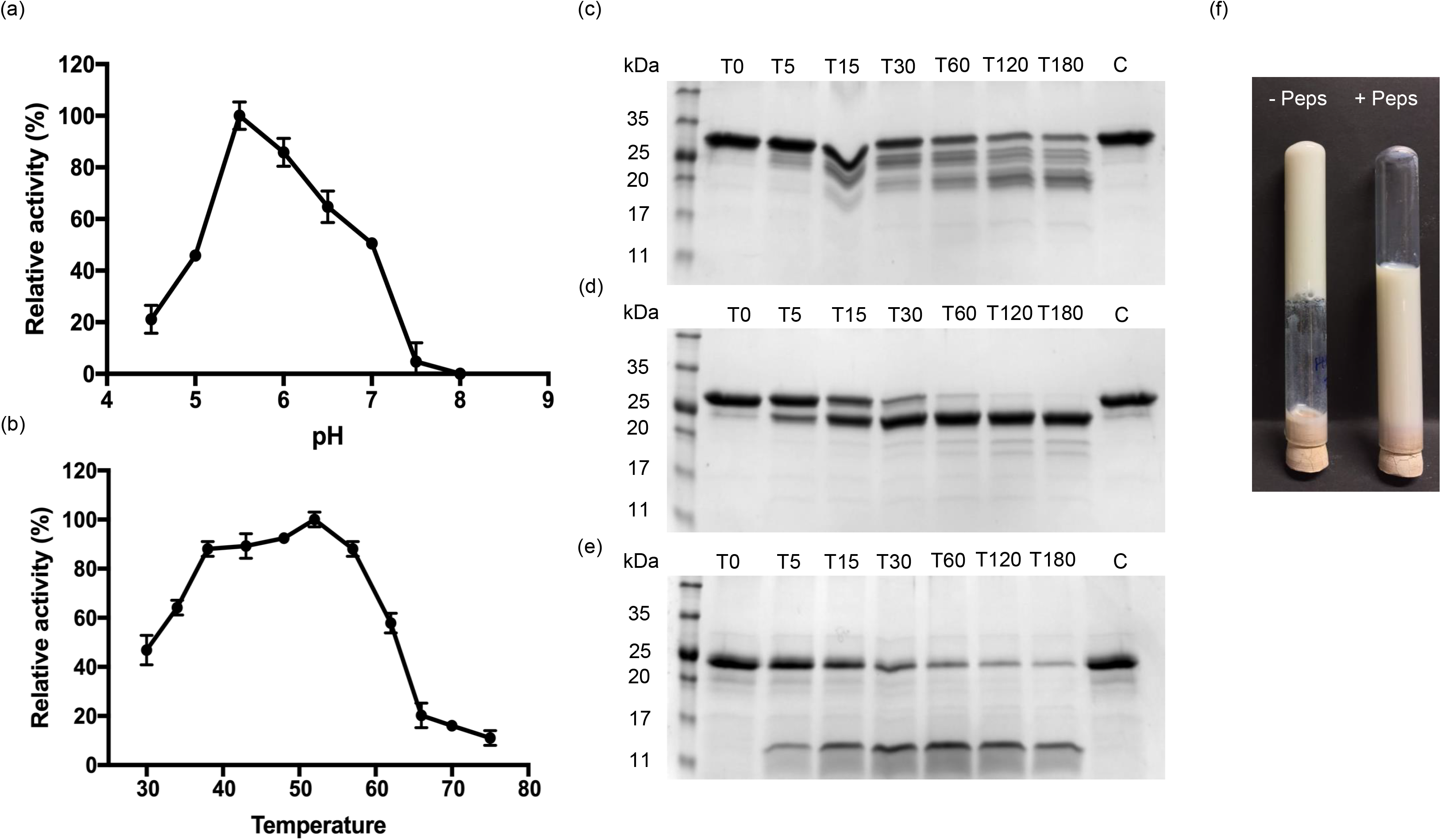
Characterization of proteolytic activity of purified cardosin B. (**a**) Determination of optimal pH (**b**) Determination of optimal temperature (**c**, **d**, **e**) SDS-PAGE analysis of α-casein, β-casein and κ-casein digestion by cardosin B, time-points are represented in minutes, C represents untreated control sample taken at T180 (**f**) Milk clotting assay using purified cardosin B with and without pepstatin A. Values are the mean of three technical replicates.

We also looked at the activity of recombinant cardosin B on milk caseins. To that end, the purified enzyme was incubated with each purified casein, α, β and κ-casein, the constituents of the casein micelle. As expected, cardosin B showed proteolytic activity over all the caseins with extensive action on α-casein (Fig. 3c). The digestion of κ-casein yielded a 13 kDa fragment (Fig. 3d) called para-κ-casein which originates from the specific cleavage of κ-casein between the Phe105-Met106 bonds ^20^. The action on α-casein was more extensive, yielding multiple bands, while that on β-casein was more specific, generating a single, 23 kDa product (Fig. 3e). These bands had previously been characterized ^21^.

The purified cardosin B was then used to determine milk clotting. The incubation of purified cardosin B with reconstituted skimmed milk led to its coagulation after 40 min, which was prevented when cardosin B was pre-incubated with pepstatin A, confirming that the effect was a result of cardosin B activity (Fig. 3f).

### CardB-dsRed fusion lines appear to delay cardosin B processing and allowed to detect an unprocessed cardosin B

Since the recombinant protein was not secreted, we generated new BY2 cell lines in which the red fluorescent protein (dsRed) was fused to the C-terminal of cardosin B. The calli expressing this construct showed a clear red signal under the fluorescence stereomicroscope, compared to the line expressing cardosin B alone (Fig. 4a).

**Figure 4.**
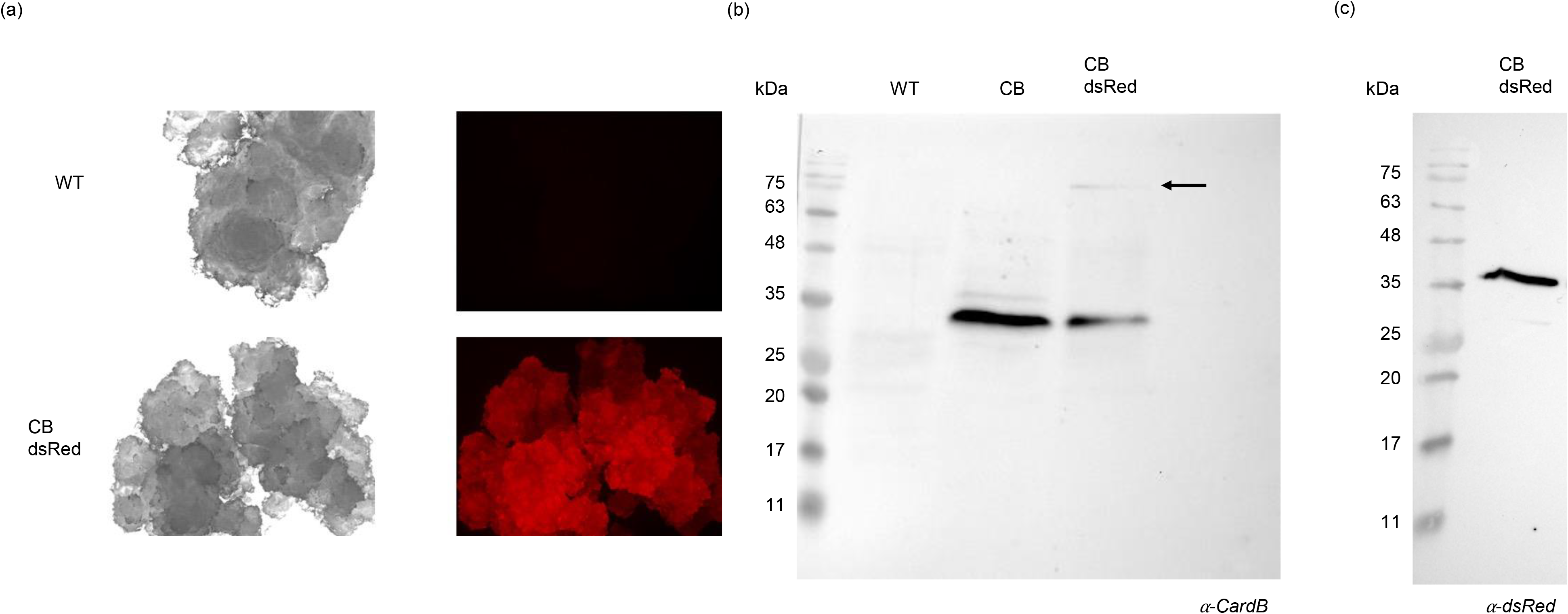
Analysis of CardB-dsRed fusion lines. (**a**) Fluorescence stereoscope visualization of WT and CardB-dsRed calli (**b**) Western blot analysis with anti-cardosin B antibody of WT, CardB and CardB-dsRed cellular extracts (**c**) Western blot analysis with anti-dsRed antibody of CardB-dsRed cellular extracts.

To further characterize this line, western blottings using antibodies against cardosin B and dsRed were performed. Anti-cardosin B antibody revealed the presence of a 34 kDa band similar to that of the lines expressing only cardosin B. However, a signal between 75 kDa and 100 kDa was also detected (Fig. 4b, arrow) which did not appear in the line expressing only cardosin B (Fig. 4b). This size is consistent with unprocessed cardosin B fused with dsRed, which may indicate a delay in cardosin B processing in this fusion cell line. Western blot analysis using an antibody against dsRed uncovered two bands, one above 35 kDa and another above 25 kDa (Fig. 4c, arrowhead). The latter may correspond to free dsRed protein with a molecular weight of 28 kDa ^22^, while the band above 35 kDa may correspond to the dsRed protein linked to the light-chain of cardosin B. To test this hypothesis, the band detected with the anti-dsRed antibody was purified by Q-sepharose and analyzed by N-terminal sequencing. The sequence obtained was SAESIV, which corresponds to the expected N-terminal sequence of the light-chain ^4^. This confirms that the dsRed signal corresponds to processed cardosin B and indicates that cardosin processing B in BY2 cells was similar that previously described for the flower ^4^.

### Activation of the unprocessed cardosin B suggests a putative role of a cysteine protease in cardosin B processing

As explained above, the size of the unprocessed cardosin B fused to dsRed conformed to that predicted. Because this protein carried a 6xHis tag, we tried to purify it using the IMAC system. Analysis by western blotting of the fractions, using the anti-cardosin B antibody, revealed two different bands corresponding to (i) the processed cardosin B in the sample plus flow-through fractions, and (ii) the unprocessed form in the eluted fraction (Fig. 5a). We, thus, obtained an enriched fraction containing the unprocessed cardosin B, which enabled a closer investigation of its processing steps.

**Figure 5.**
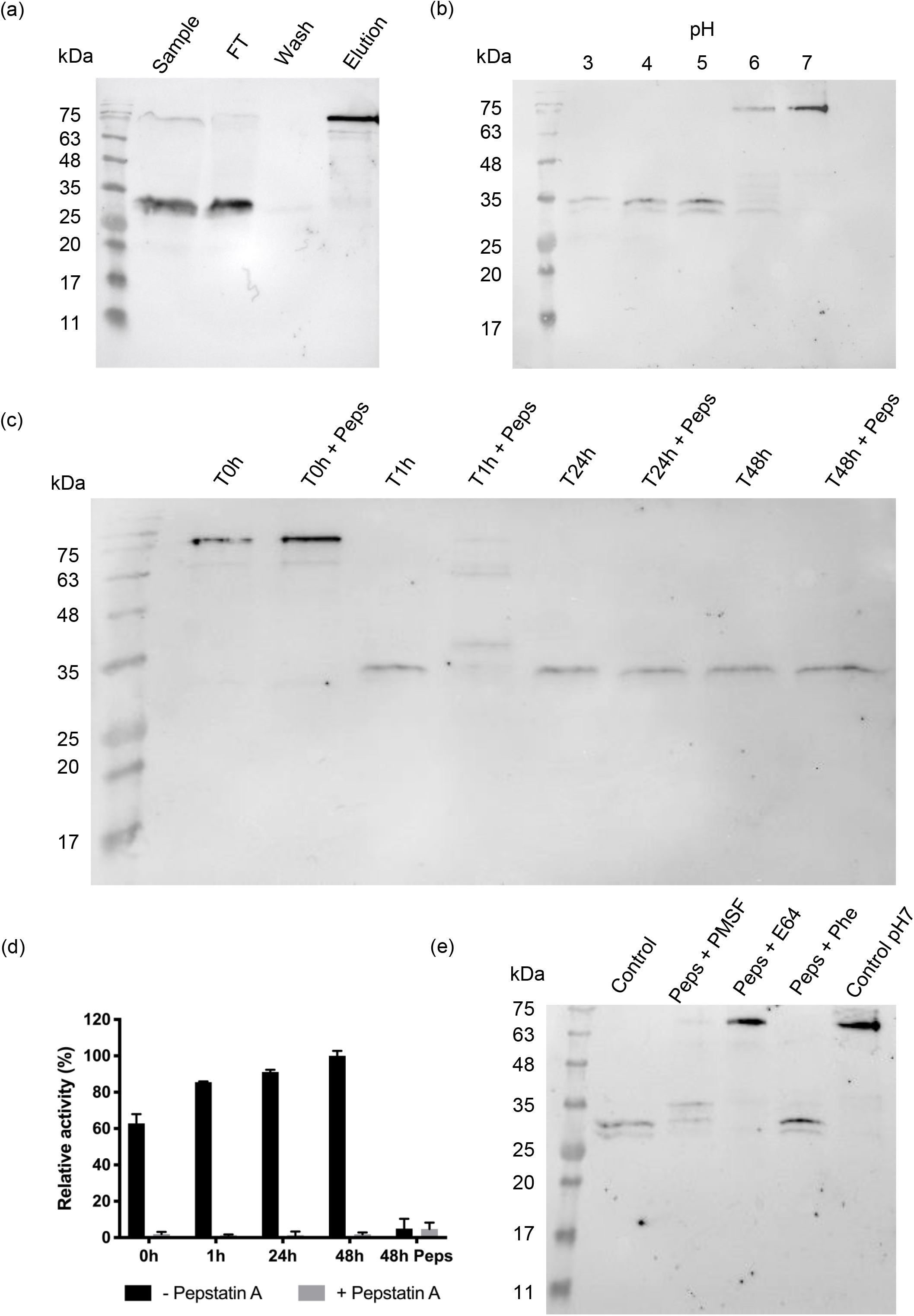
CardB-dsRed fractionation by IMAC and activation assays of unprocessed cardosin B (**a**) Western blot analysis of the fractionation of cardB-dsRed cell extract by IMAC with anti-cardosin B antibody (**b**) Western blot analysis of the effect of pH on unprocessed cardosin B (**c**) Western blot analysis of the effect of pepstatin A on cardosin B at pH 4. Time-points of samples with and without pepstatin A were collected at 0h, 1h, 24h and 48h after incubation at pH 4 (**d**) Activity assay of samples with and without pepstatin A (**e**) Western blot analysis of the combined effect of pepstain A with other protease inhibitors on cardosin B processing. Peps, pepstatin A; PMSF, phenylmethylsulfonyl fluoride; Phe, phenanthroline.

Cardosin A has been described as being activated after incubation at pH 4 in a process inhibited by pepstatin A, a specific AP inhibitor, which suggests that cardosin A is self-activating ^11^. To find out if the same occurs for cardosin B, we incubated fractions containing the unprocessed enzyme at various pH levels and used western blotting to assess the protein patterns. We observed that acidification reduces the size of the detected band from 75 kDa to 34 kDa (Fig. 5b). To understand if this shift was caused by the activity of cardosin B, samples were incubated with pepstatin A prior to incubation at pH 4. The presence of pepstatin A delayed the band shift to 34 kDa but did not lead to complete inhibition (Fig. 5c). To determine if this observation might be explained by ineffective pepstatin A inhibition, samples at different incubation time-points, with and without the compound, were collected and used in an activity assay. Surprisingly, the proteolytic activity in the pepstatin A treated samples was completely blocked compared to those without pepstatin A pre-treatment, which indicated that cardosin B was in fact inhibited. To further confirm these results, both samples collected at time-point 48 h (with and without pre-treatment with pepstatin A) were treated with pepstatin A prior to the activity assay. In this assay, reduced activity confirmed that the proteolytic activity measured in the non-treated samples was mostly due to cardosin B (Fig. 5d). Furthermore, the single use of protease-specific inhibitors did not prevent processing (Fig. S4), which suggests that it may involve additional classes of proteases. We then combined pepstatin A with inhibitors of other protease classes. The combined effect of pepstatin A and E64 was able to prevent cardosin B processing, indicating the putative role of cysteine proteases in cardosin B processing (Fig. 5e).

In order to identify these cysteine proteases, a proteome characterization was performed in the eluted fraction containing unprocessed cardosin B, and the results were compared to those of the *Nicotiana tabacum* database. Three cysteine proteases were identified (Table 1). CP15 shares 97% identity with low-temperature induced cysteine protease and 76.6% with CP6. CP6 shares 77% identity with low-temperature induced cysteine protease. Interestingly, all the proteases uncovered were papain-like cysteine proteases, with a granulin-domain at the C-terminal whose function is still unknown ^23–25^.

**Table 1:** Cysteine proteases detected in the eluted fraction containing unprocessed cardosin B by mass spectrometry.

| Uniprot accession number | Protein name |
| --- | --- |
| tr\|T2BQ67\|T2BQ67_TOBAC | Cysteine protease CP15 |
| tr\|T2BRU8\|T2BRU8_TOBAC | Cysteine protease CP6 |
| tr\|A0A1S4B1Z6\|A0A1S4B1Z6_TOBAC | Low-temperature induced cysteine protease |

## Discussion

Significant research over the years has looked into systems for expressing APs from cardoon for potential industrial application; however, a viable system applicable to all APs has yet to be developed. We assessed the use of BY2 cells as a production platform for these APs, using cardosin B as a model.

In BY2 cells expressing recombinant cardosin B, the protein was only detected in the cellular extracts and not in the spent medium. Moreover, the protein was found only in its processed form, and no intermediary forms were identified. Similar results have been reported for cardosin B in tobacco and *Arabidopsis thaliana* ^12^. This contrasts with cardosin A, for which intermediary forms of the protein were detected ^26^. The detection of only the processed form suggests a fast processing of the enzyme. This rapid processing was also described by other authors, which argued that maturation of cardosin B must occur as early as the Endoplasmic Reticulum-Golgi Apparatus. However, the precise location of activation remains unknown ^12^. An Endo H assay indicated that cardosin B produced in BY2 cells contains high-mannose glycosylation, which is consistent with the pattern found in other heterologous systems as well as cardoon flower ^12^. This suggests that cardosin B processing in differentiated systems may also occur in undifferentiated systems such as BY2 cells.

Cardosin B purification from BY2 cells yielded a fraction with pure cardosin B. It presented the expected 34 kDa heavy-chain as well as a lower band corresponding to the light-chain; however, the latter had a smaller size than that of the light-chain cardosin B from the cardoon flower ^4^. MS analysis of each band showed that the protein was not subjected to any other proteolytic event than that required to process cardosin B, thus suggesting that the size of the cardosin B light-chain may in fact be smaller than previously reported.

The purified enzyme showed optimal activity at pH 5.5 and 50 °C. The proteolytic activity of cardosin B on cow milk caseins has been studied and shown to have a broader specificity than cardosin A ^27^. We found that purified cardosin B showed the predicted specific activity on κ-casein and a broader proteolytic effect on α-and β-caseins. The purified cardosin B was also able to clot milk, supporting its ability to be used in cheese manufacturing. Taken together, these results suggest that BY2 cells may represent a viable alternative source of cardosins. Nevertheless, the low yields of purified protein still hinder industrial scale application. Pure enzymes were effectively produced, with however high levels of protein loss to precipitation during acidic cellular extraction. As such, this procedure requires a tradeoff between purity and quantity. The use of a 6xHis tag at the C-terminal proved to be of limited usefulness for improving purification. Future research may look into other strategies, such as manipulating the trafficking pathway towards cardosin B secretion to the culture medium.

The dsRed fusion line shined light on cardosin B processing in BY2 cells. Anti-dsRed antibody detected a band higher than 35 kDa. The subsequent N-terminal analysis of this band revealed the expected amino acid sequence of the light-chain. The fact that anti-dsRed primarily detected this form points, once again, to early cardosin B processing, which appears not to have suffered impairment by the dsRed fusion. However, the appearance of a band between 75 kDa and 100 kDa, which was not found in the cardosin B expressing lines, suggests that dsRed fusion may delay cardosin B processing thereby leading to its detection. The size is consistent with that predicted for unprocessed cardosin B fused to dsRed. This delay may be related to dsRed being an obligate tetramer ^22^, which means that the four subunits must be linked. The size of this complex may interfere with normal cellular trafficking, leading to its detection.

Little is known about cardosin B processing. The enriched fraction containing unprocessed cardosin B allowed us to further investigate its activation mechanism. As expected, the acidic pH led to a size change in the detected heavy-chain consistent with cardosin processing ^28^. The same has already been reported for recombinant cardosin A produced in *E. coli* ^11^. In this study, pepstatin A inhibited the process, leading the authors to argue that cardosin A may be self-activating ^11^. We assessed whether the same was true for cardosin B and found that pepstatin A was unable to fully inhibit cardosin B processing. The use of protease-specific inhibitors also failed to prevent it, which suggests that more than one protease class is involved. Combining pepstatin A with other protease inhibitors suggested the putative role of a cysteine protease in cardosin B processing and proteome analysis identified three potential candidates. Interestingly, all cysteine proteases identified by MS contained a granulin-domain at their C-terminus. Although its role is still unknown, the fact that the granulin-domains were uncovered may help explain a possible interaction between the cysteine proteases and cardosin B. In human cells, progranulin has been found to interact with both prosaposin ^29^, a glycoprotein involved in the lysosomal trafficking pathway, and cathepsin D, a human aspartic protease ^32,33^). Like most typical plant APs, cardosin B contains a saposin-like domain corresponding to the PSI segment. Thus, potential interaction between the cysteine proteases containing a granulin-domain and cardosin B may prove similar to that observed in human cells, although in plants the saposin-domain and the aspartic protease are part of the same enzyme. This putative interaction may further explain the rapid processing of cardosin B as opposed to cardosin A and increase the range of possible mechanisms behind plant AP regulation and processing. A preliminary observation of the cardB-dsRed cells under a confocal microscope seemed to indicate that the red signal covered most of the cell volume, which may be compatible with sorting to the vacuole and thus following the same pathway described in other heterologous expression systems ^12^. Additionally, in this line a cluster of vesicles was observed at the nuclear periphery that was absent in WT cells and which may be ER-derived ^32^. Although the study of the trafficking pathway of the recombinant protein was beyond the scope of this investigation, these observations open avenues for further research and highlight the importance of understanding the fundamental aspects of the plant secretory pathway so as to fully realize their biotechnological potential.

The use of plant cells as a platform proved viable and able to produce active cardosin B enzyme, which represents an improvement relative to other production platforms such as microbial systems. The activation and trafficking of plant APs seem to be intertwined and closely regulated at the cellular level. Plant-based platforms offer the same structural and basal mechanisms that occur in nature for the expression and processing of APs, thus making their production similar to that occurring in the flower. This offers a new perspective on the biotechnological use of cardoon APs that may contribute to making plant-based platforms the solution of choice for future industrial cardoon AP applications.

## Methods

### Plasmid constructs

Cardosin B coding sequence (EMBL no. AJ237674) without the native signal peptide was synthesized by NZYTech (NZYTech, Portugal) with the addition of NcoI and SalI restriction sites at the 5’and 3’ends respectively. The gene was then cloned into the pTRA vector (kindly offered by Thomas Rademacher, Aachen, Germany) using the NcoI and SalI sites. The resulting vector contained the cardosin B coding sequence controlled by the enhanced Cauliflower Mosaic Virus 35SS constitutive promoter, followed by a 5’UTR from Chalcone Synthase (CHS) and a murine signal peptide (LPH). The vector also contains a 6xhistidine tag used for purification purposes at the C-terminus and a 35S terminator (Fig. S5a). A second vector was built with a fusion of cardosin B to dsRed. The dsRed sequence was amplified from the binary plasmid pLB52 ^33^ with primers containing SalI and NotI restriction sites at the 5’and 3’respectively. The dsRed gene was then cloned into pTRA-CardB vector with SalI and NotI, resulting in a vector denominated pTRA-CardB-dsRed (Fig. S5b).

### Agrobacterium transformation of tobacco BY2 cells

Both constructs described above were transformed into *Agrobacterium tumefaciens* strain GV3101::pMP90RK by the freeze-thaw method. BY2 cells were transformed with each construct by *A. tumefaciens* as described ^34^ with slight modifications. Two days after co-culture, the cells were transferred to gelrite plates containing kanamycin (100mg/L) (NZYTech, Portugal) and ticarcillin disodium/clavulanate potassium (500mg/L) (Duchefa, Netherlands) to select transformants and eliminate Agrobacterium. Growing microcalli were isolated and subcultured to fresh plates containing the same antibiotics, every two weeks. Ticarcillin disodium/clavulanate potassium concentration was decreased to half in each subculture until complete removal. Liquid cultures were established from growing callus.

### Protein extractions and spent medium

At day 7 of growth, BY2 cultures were collected and paper-filtered to separate cells from the spent medium. Cells were macerated in liquid nitrogen using a mortar and pestle and then homogenized in 100 mM Tris-HCl, pH 7 in a 1:1 ratio (w/v). The extracts were centrifuged at 16000 g for 15 min at 4 °C and the supernatant was kept. The spent medium was collected and centrifuged at 16000 g for 15 min at 4 °C to remove cell debris. For the optimization of buffer pH for cell extraction, BY2 cell extracts were prepared using buffers with various pH values: 100 mM citrate buffer (pH 3-5), 100 mM phosphate buffer (pH 6) and 100 mM Tris-HCl (pH 7-8). Total soluble protein content of the protein extracts was determined using Bradford protein assay (Bio-Rad, USA) with bovine serum albumin (BSA) as standard, following manufacturer’s instructions. Cardosin B content was determined using FIJI software based on the relative intensity of bands detected on the western blot.

### Western blotting analyses

Cellular protein extracts and/or spent medium were loaded onto a 12.5% polyacrylamide gel for SDS-PAGE analysis. After electrophoresis, the proteins were transferred to a nitrocellulose membrane using Trans-blot turbo (Bio-Rad, USA). The membrane was incubated in 5% (w/v) skimmed milk and 3% (w/v) BSA (NZYTech, Portugal) in phosphate buffered saline containing 0.1% (v/v) Tween 20 (PBS-T) for 1 h at room temperature (RT). The membrane was incubated with an anti-cardosin B antibody (Antibody generated by HuCAL technology, Bio-Rad, USA) as described in ^16^ diluted in PBS-T (0.4 μg/ml) or anti-dsRed (sc-3909098, Santa-Cruz Biotech, USA) diluted 1:200 in PBS-T, for 1 h at RT and then at 4 °C overnight. Anti-human IgG peroxidase conjugated (A00166, GenScript, USA) or anti-mouse IgG peroxidase conjugated (A4416, Sigma Aldrich, USA) were used as secondary antibodies at dilutions 1:5000 and 1:4000 in PBS-T, respectively, under agitation for 2 h at RT. The signal was detected using Hyperfilm ECL (GE healthcare biosciences, USA) or Supersignal™ West femto maximum sensitivity substrate (Thermo Fisher Scientific, USA) in a ChemiDoc XRS+ system (Bio-Rad, USA). Protein molecular weight marker was NZYColour Marker II (NZYTech, Portugal) or Precision Plus Protein WesternC Standards (Bio-Rad, USA).

### Endoglycosidase digestion assay

Endoglycosidase digestion was performed according to ^37^ with a few modifications. Cell extracts from BY2 lines transformed with pTRA-CardB were prepared as previously described. Then, 3 μL of 10X concentrated denaturing buffer (500 mM sodium citrate, pH 5.5, 2% SDS, 10% β-mercaptoethanol) was added to 30 μL of cell extract and samples were boiled at 100 °C for 10 min. After boiling, samples were incubated for 5 min on ice and 3 μL of 10X G5 buffer (New England BioLabs, UK) were added to the samples together with 1.5 μL of 25X concentrated protease inhibitor cocktail (Roche, USA) and endoglycosidase H (2.5 mU) (Roche, USA). Samples were incubated at 37 °C overnight. Control samples were treated without endoglycosidase H. Finally, 0.25 volumes of 4X SDS-PAGE loading buffer was added to the samples and analyzed by SDS-PAGE and western blot as previously described.

### Purification of cardosin B and cardosin B-dsRed

Cellular extracts from BY2 cell lines producing cardosin B were prepared as mentioned above. The macerated cells, approximately 100 g, were homogenized in 100 mM citrate buffer, pH 3 in a 1:1 ratio (w/v), and then concentrated with an Amicon Ultra-15 centrifugal filter (Merck Millipore, Germany). The concentrated sample was subjected to buffer exchange by dialysis against 20mM Tris-HCl pH7,4.

Purification of cardosin B was performed and monitored by chromatography using an AKTA system (GE Healthcare, USA). 20X concentrated protein extract (10 ml) was loaded in an anionic-exchange Q-sepharose column (GE Healthcare, USA) previously equilibrated with 20 mM Tris-HCl pH 7.4. Washing was performed using the same buffer. Elution was performed using a gradient ranging from 0 to 100% with 1 M NaCl in the same buffer. Each eluted fraction was collected in 10 mL and analyzed by SDS-PAGE and western blot with anti-cardosin B antibody as described previously. Positive fractions for cardosin B were pooled and concentrated.

For cardosin B-dsRed fusion lines, eluted fractions were observed under a Zeiss Stereo Microscope Axio Zoom V16 with a filter set 63 HE mRFP (EX BP 572/25 BS FT 590, EM BP 629/62).

Images were acquired with a Zeiss Axiocam 503 mono camera and processed with Adobe Photoshop. Fractions presenting fluorescence were pooled and analyzed by SDS-PAGE and western blotting. Purification of the unprocessed form of cardosin B was performed by IMAC using a 1 mL HisTrap column (GE Healthcare, USA) following manufacturer’s instructions. Elution was performed with an imidazole gradient of 100 to 500 mM. All fractions were analyzed by SDS-PAGE and western blot with anti-cardosin B antibody.

### Determination of optimal pH and temperature for maximum activity of purified cardosin B

Optimal pH for cardosin B activity was assayed using azocasein (Sigma, USA) as substrate. 1 μL of purified cardosin B (11 μg/mL) was added to 150 μL of 0.5% azocasein solution (w/v) ranging various pH values, using 100 mM citrate buffer (pH 4.-5.5), 100 mM phosphate buffer (pH 6-6.5) or 100 mM Tris-HCl (pH 7-8). Samples were incubated at 37 °C for 1 h. Optimum temperature was assessed by incubating the samples in 100 mM citrate buffer pH 5.5 at various temperatures ranging from 35 °C to 70 °C for 1 h. The reaction was stopped by the addition of 50 μL of 10% (w/v) trichloroacetic acid (TCA). Samples were incubated 30 min on ice, followed by centrifugation at 10000 g for 10 min. 100 μL of supernatant were added to 100 μL of 1 N NaOH and the absorbance of the solution was measured at 440 nm using an Epoch microplate spectrophotometer (BioTek, USA).

### Caseins hydrolysis assay

Purified cardosin B (11 μg/mL) was added to pure α-casein, β-casein and κ-casein (Sigma,USA) (200 μg/mL) in 100 mM sodium phosphate buffer pH 6.4 in a ratio 1:100 (v/v) and incubated at 37 °C. Time-points 0, 5, 15, 30, 60, 120 and 180 min were collected and analyzed. Control for non-enzymatic digestion was performed using 20 mM Tris-HCl pH 7.4 instead of cardosin B and was collected at 180 min.

### Milk clotting assay

100 μL of purified cardosin B was incubated with pepstatin A (4 μM) for 30 min at RT while the control sample was incubated with methanol instead of pepstatin A. After 30 min, samples were added to 5 mL of reconstituted skimmed milk (12% w/v in water, 10 mM CaCl_2_, pH 6.4) and incubated at 30 °C in a water bath. Milk clotting was assessed by visual observation by manually inverting the tubes.

### N-terminal analysis

Protein bands were run in a 15% polyacrylamide gel and transferred to a PVDF membrane using a mini trans-blot cell (Bio-Rad, USA). The membrane was stained with Coomassie blue R-250 and the bands were cut with a sterilized scalpel. Protein bands were identified using the Edman reaction and a Procise 491 HT Protein Sequencer to determine the N-terminal sequences (Analytical Laboratory, Analytical Services Unit, ITQB NOVA).

### Study of the activation and inhibition of the unprocessed form of Cardosins B

Purified fractions containing the unprocessed form of cardosin B were concentrated 10X using amicon ultra-15 (3kDa MWCO) (Millipore, USA). For cardosin B processing assay, concentrated fractions were diluted 1:1 in phosphate buffer 100 mM (pH 7 and 6) or citrate buffer 100 mM (pH 5, 4 and 3) and incubated at 37 °C, samples were collected and analyzed by western blot. For the inhibition assay, the concentrated fraction was incubated with pepstatin A (4 μM) for 30 min at RT before dilution in citrate buffer 100 mM pH 4, the control sample was incubated with methanol. Time-points were collected at 0 h, 1 h, 24 h and 48 h. The time-points collected from the inhibition assay were also used to assess activity using azocasein as substrate as described above. For double inhibition, the concentrated fraction was incubated with pepstatin A and independently combined with phenylmethylsulfonyl fluoride (PMSF) (1 mM), E64 (100 μM) or Phenanthroline (10 mM). Samples were incubated at 37 °C for 24 h, collected and analyzed by western blot.

### Mass spectrometry analysis of purified cardosin B and unprocessed cardosin B fraction

The three bands of purified cardosin B from BY2 cells and the eluted fraction containing the unprocessed form of cardosin B were used for protein/proteome characterization. Both samples were loaded in a 10% SDS-PAGE gel. The sample with the eluted fraction containing the unprocessed form run for 15 min and the sample with purified cardosin B was run until the three bands were clearly separated. Afterward, the gel was stained with BlueSafe (NZYTech, Portugal) and protein bands were cut out using a scalpel blade. Protein identification was performed by mass spectrometry as previously described ^35^, with exception of the Uniprot database used for protein identification. Gel bands were reduced in 10 mM DTT (Sigma) for 40 min at 56⁰C, and alkylated in 55 mM iodoacetamide (Sigma) for 30 min in the dark. Excessive iodoacetamide was quenched by further incubation with DTT (10 mM for 10 min in the dark). The resulting sample was digested overnight with trypsin (Promega, Madison, WI, USA) at 37⁰C (1:50 protein/trypsin ratio) and cleaned up with a C18 column (OMIX C18, Agilent, Santa Clara, CA, USA). Nano-liquid chromatography-tandem mass spectrometry (nanoLC-MS/MS) analysis was performed on an ekspert™ NanoLC 425 cHiPLC^®^ system coupled with a TripleTOF^®^ 6600 with a NanoSpray^®^ III source (Sciex). Peptides were separated through reversed-phase chromatography (RP-LC) in a trap-and-elute mode. Trapping was performed at 2 μl/min on a NanoLC Trap column (Sciex 350 μm x 0.5 mm, ChromXP C18-CL, 3 μm, 120 Å) with 100% A for 10 min. The separation was performed at 300 nl/min, on a NanoLC column (Sciex 75 μm x 15 cm, ChromXP 3C18-CL-120, 3 μm, 120 Å). The gradient was as follows: 0-1 min, 5% B (0.1% formic acid in acetonitrile, Fisher Chemicals, Geel, Belgium); 1-46 min, 5-35% B; 46-48 min, 35-80% B; 48-54 min, 80% B; 54-57 min, 80-5% B; 57-75 min, 5% B. Peptides were sprayed into the MS through an uncoated fused-silica PicoTip™ emitter (360 μm O.D., 20 μm I.D., 10 ± 1.0 μm tip I.D., New Objective, Oullins, France). The source parameters were set as follows: 15 GS1, 0 GS2, 30 CUR, 2.5 keV ISVF and 100 ⁰C IHT. An information dependent acquisition (IDA) method was set with a TOF-MS survey scan of 400-2000 m/z. The 50 most intense precursors were selected for subsequent fragmentation and the MS/MS were acquired in high sensitivity mode for 40 msec. The obtained spectra were processed and analyzed using ProteinPilot™ software, with the Paragon search engine (version 5.0, Sciex). A UniProt database (76,177 entries, accessed in 12/07/2020) containing the sequences of the *Nicotiana tabacum* was used for the analysis of the fraction containing the unprocessed form of cardosin B. For the analysis of purified cardosin B bands, the expected amino acid sequence of recombinant cardosin B was added to the database. The following search parameters were set: iodoacetamide, as Cys alkylation; trypsin, as digestion; TripleTOF 6600, as the Instrument; gel-based ID as Special factos; ID focus as biological modifications and amino acid substitutions; search effort as thorough; and an FDR analysis. Only the proteins with unused protein score above 1.3 and 95% confidence were considered.

## Acknowledgements

This work was supported by Fundação para a Ciência e Tecnologia (FCT, Portugal) through project ref. PTDC/BAA-AGR/30447/2017, Research Unit ref. UIDB/04551/2020 and PhD Fellowship to AF (ref. PD/BD/114488/2016, Plants for Life PhD Program). MS data were obtained by the UniMS – Mass Spectrometry Unit, ITQB/IBET, Oeiras, Portugal. The authors wish to thank Bruno Alexandre for help with the MS analysis, and Andreas Schiermeyer and Catarina Pimentel for fruitful discussions and critical reading of the manuscript.

## Author contributions

AF and RA conceived and designed the work. AF performed the experiments. AF and RA analyzed the data and wrote the manuscript.

