## Supplementary Material for "Expression of recombinant cardosin B in tobacco BY2 cells: an alternative system for the production of active milk clotting enzymes"

**Figure S1**

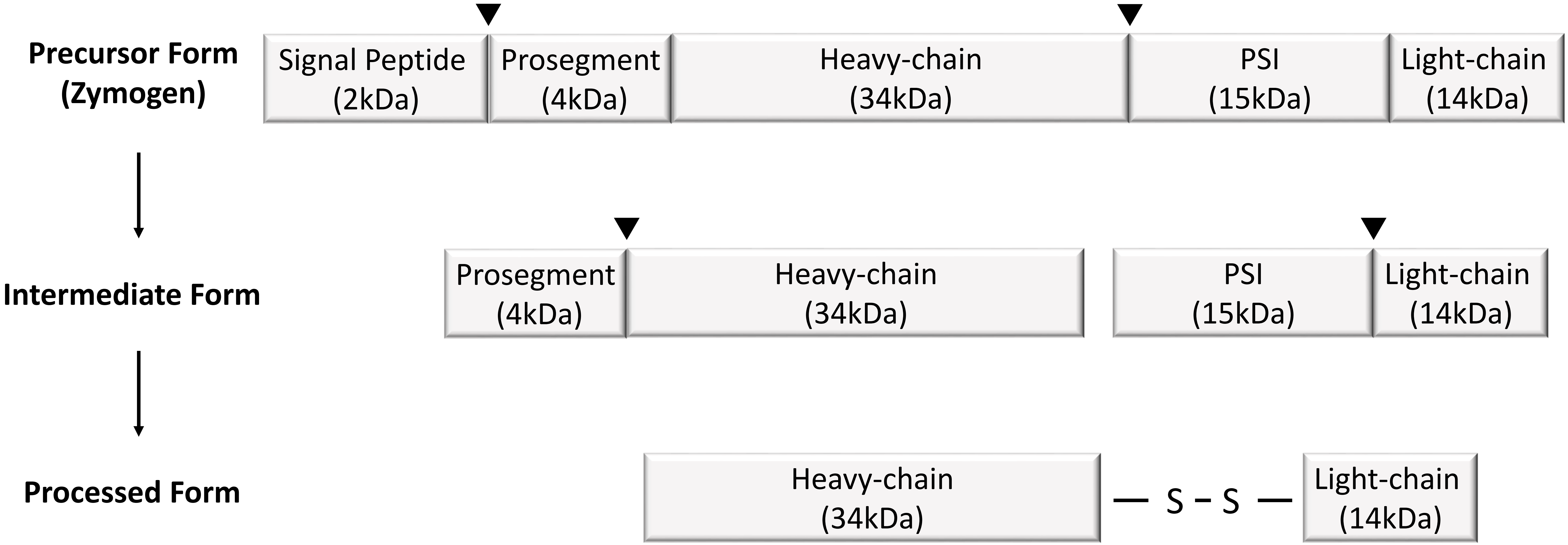


**Figure S1.** Schematic representation of cardosin B processing steps. Arrowheads indicate cleavage site during processing. PSI – Plant Specific Insert. S-S indicates disulfide bond between heavy- and light-chains.

**Figure S2**

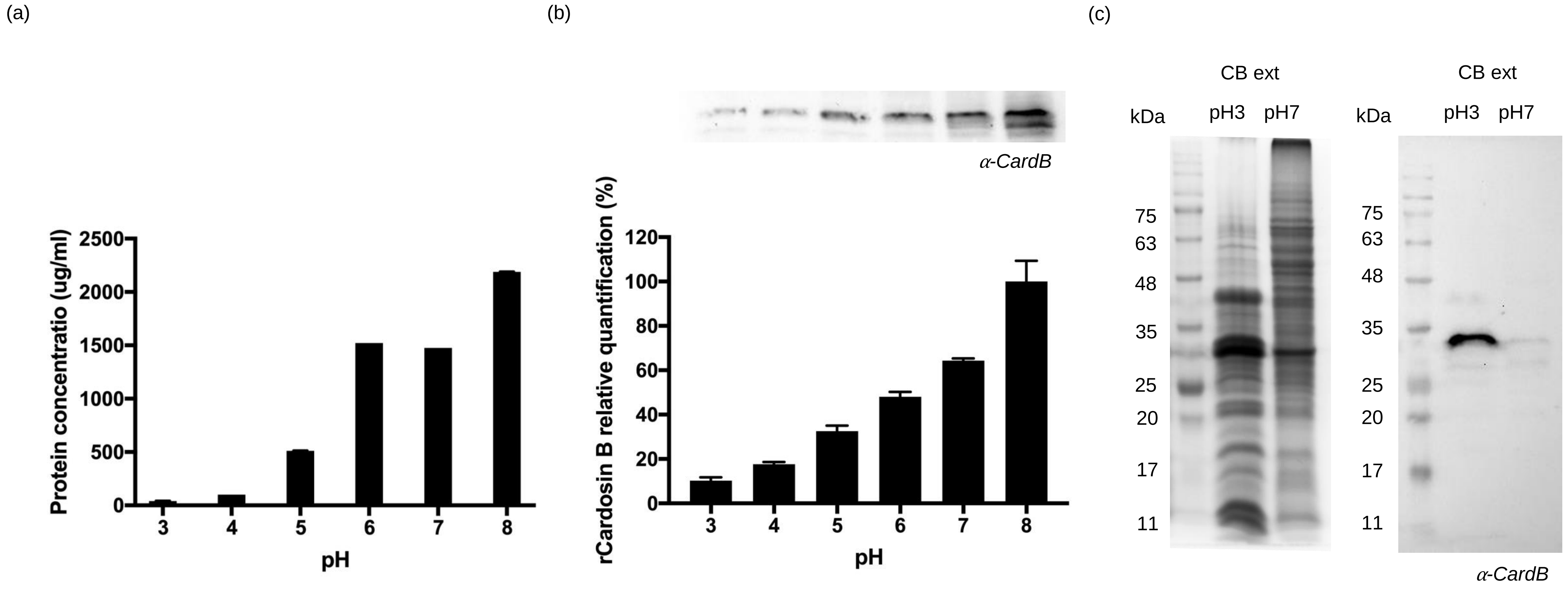


**Figure S2.** Optimization of cellular extractions. (a) Effect of pH on cell extract protein concentration (b) Effect of pH on cardosin B extraction and western blot analysis of the protein extracts at various pH values. The values are the mean of three western blot replicates (c) SDS-PAGE and western blot analysis of cell extracts at distinct pH values leaded with the same protein concentration.

**Figure S3**

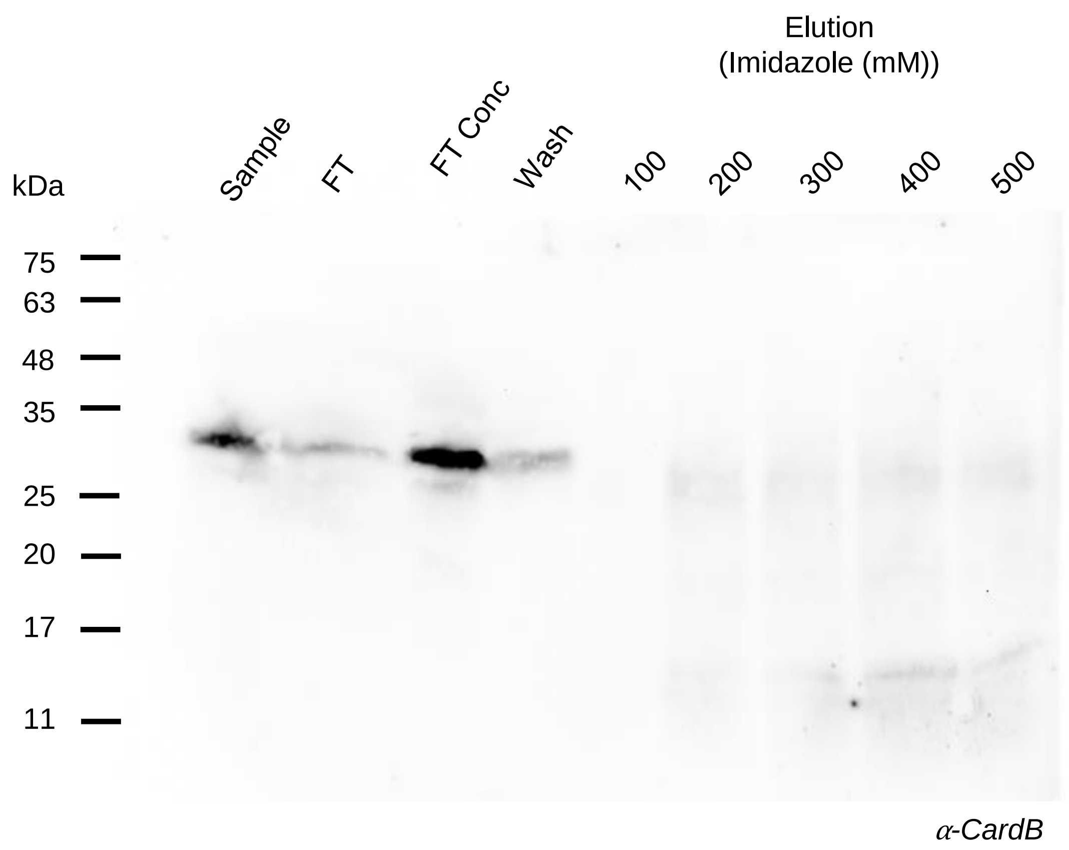


**Figure S3.** Western blot analysis of the purification of cardosin B by immobilized metal affinity using a His Trap™ column.

**Figure S4**

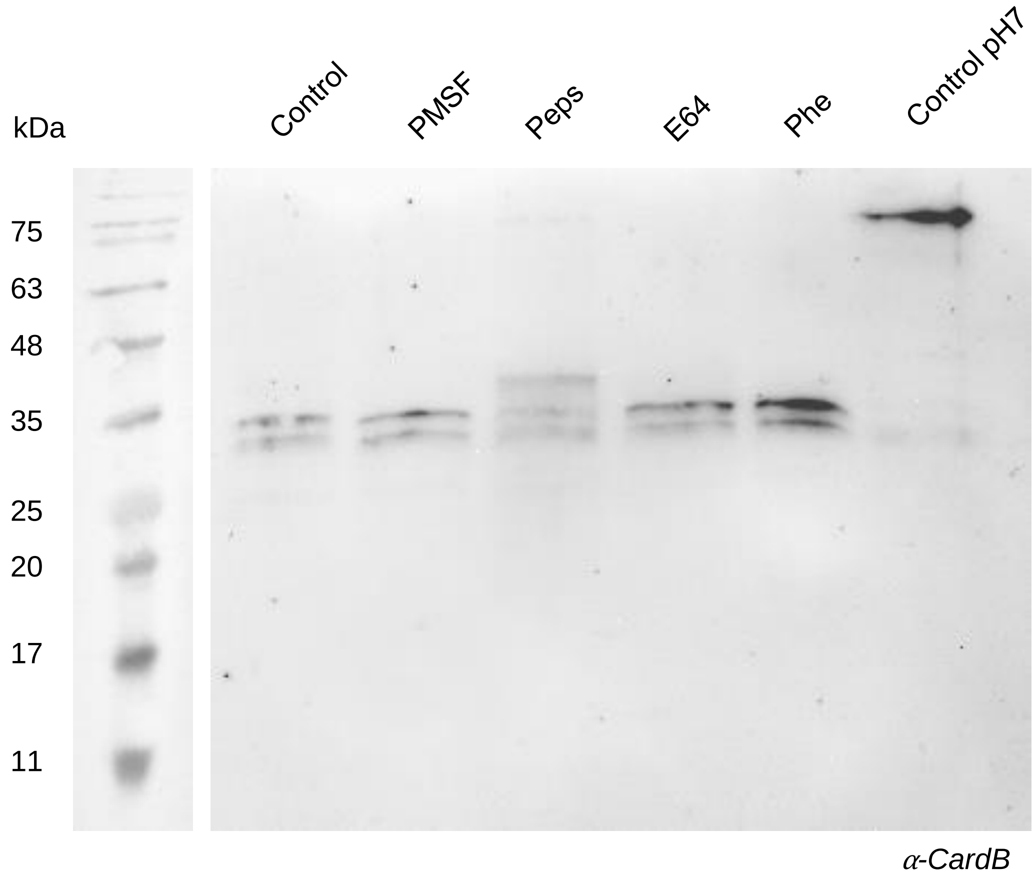


**Figure S4.** Western blot analysis of activation assay of unprocessed cardosin B with protease class specific inhibitors. Peps, pepstatin A; PMSF, phenylmethylsulfonyl fluoride; Phe, phenanthroline.

**Figure S5**

(a)

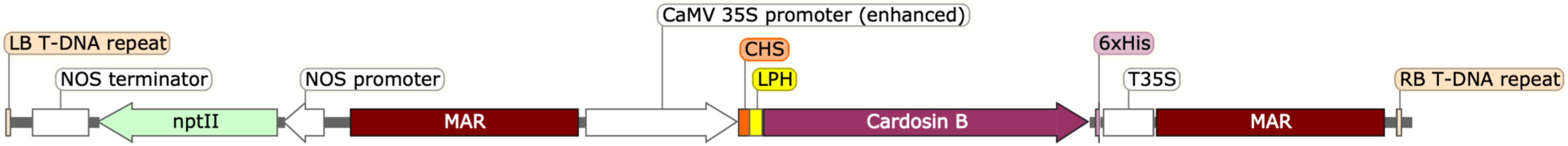


(b)

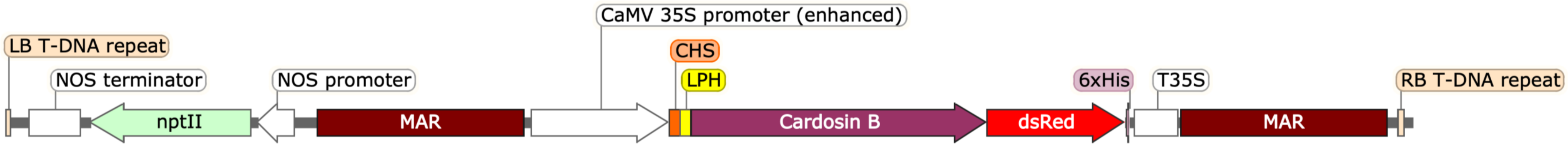


**Figure S5.** Schematic representation of the T-DNA cassette used in each vector. (a) pTRA-CardosinB (b) pTRA-CardosinB-dsRed. LB, Left border; nptII, kanamycin resistance marker; MAR, matrix attachment region; CaMV 35SS promoter, cauliflower mosaic virus 35S promoter; CHS, 5´UTR from chalcone synthase; LPH, murine signal peptide; 6xHis, six histidine tag; T35S, 35S terminator; RB, Right border.
